# A Framework to Simulate Cortical microtubule Dynamics in Arbitrary Shaped Plant Cells

**DOI:** 10.1101/169797

**Authors:** Bandan Chakrabortty, Ben Scheres, Bela Mulder

## Abstract

Plant morphogenesis is strongly dependent on the directional growth and the subsequent oriented division of individual cells. It has been shown that the plant cortical microtubule array plays a key role in controlling both these processes. This ordered structure emerges as the collective result of stochastic interactions between large numbers of dynamic microtubules. To elucidate this complex self-organization process a number of analytical and computational approaches to study the dynamics of cortical microtubules have been proposed. To date, however, these models have been restricted to 2D planes or geometrically simple surfaces in 3D, which strongly limits their applicability as plant cells display a wide variety of shapes. This limitation is even more acute, as both local as well as global geometrical features of cells are expected to influence the overall organization of the array. Here we describe a framework for efficiently simulating microtubule dynamics on triangulated approximations of arbitrary three dimensional surfaces. This allows the study of microtubule array organization on realistic cell surfaces obtained by segmentation of microscopic images. We validate the framework against expected or known results for the spherical and cubical geometry. We then use it to systematically study the individual contributions of global geometry, edge-induced catastrophes and cell face-induced stability to array organization in a cuboidal geometry. Finally, we apply our framework to analyze the highly non-trivial geometry of leaf pavement cells of *Nicotiana benthamiana* and *Hedera helix.* We show that our simulations can predict multiple features of the array structure in these cells, revealing, among others, strong constraints on the orientation of division planes.

## Introduction

It is well known that the cortical microtubule (hereafter abbreviated to MT) cytoskeleton in plant cells plays a decisive role in controlled cell expansion and oriented cell division, which together drive the plant morphogenesis^1^. In the absence of MT organizing centers like centrosomes ^2^, higher plants establish an ordered array of MTs at the cell cortex, the so-called cortical array (CA)^3^. Recent studies have revealed that cell shape may have influence on the orientation of the CA^4^. On the other hand, the orientation of the CA controls cell expansion and cell anisotropy, by guiding the deposition of cellulose synthase complexes along the MTs^5–8^. Through this coupling, the CA in turn can influence the cell shape, essentially setting up a morphogenetic feedback loop. This loop is possibly also amplified by a mechanical feedback mechanism discussed in^9^. Therefore, understanding both anisotropic cell expansion and oriented cell division, requires understanding the formation of the ordered CA from an initially disordered state just after cell division^10^.

For non-growing cells, the influence of cell shape on the formation of the CA will be static in nature and thus determined by geometrical features alone. As illustrated in Fig 1, a significant variety in these features is observed between different cell types. For growing cells, the evolving cell shape may generate a corresponding dynamic influence on the CA formation process. If growth is slow compared the collective dynamical timescale of the MTs, a quasi-static approach which samples cell shapes at different time points may still be a reasonable approximation.

**Figure 1.**
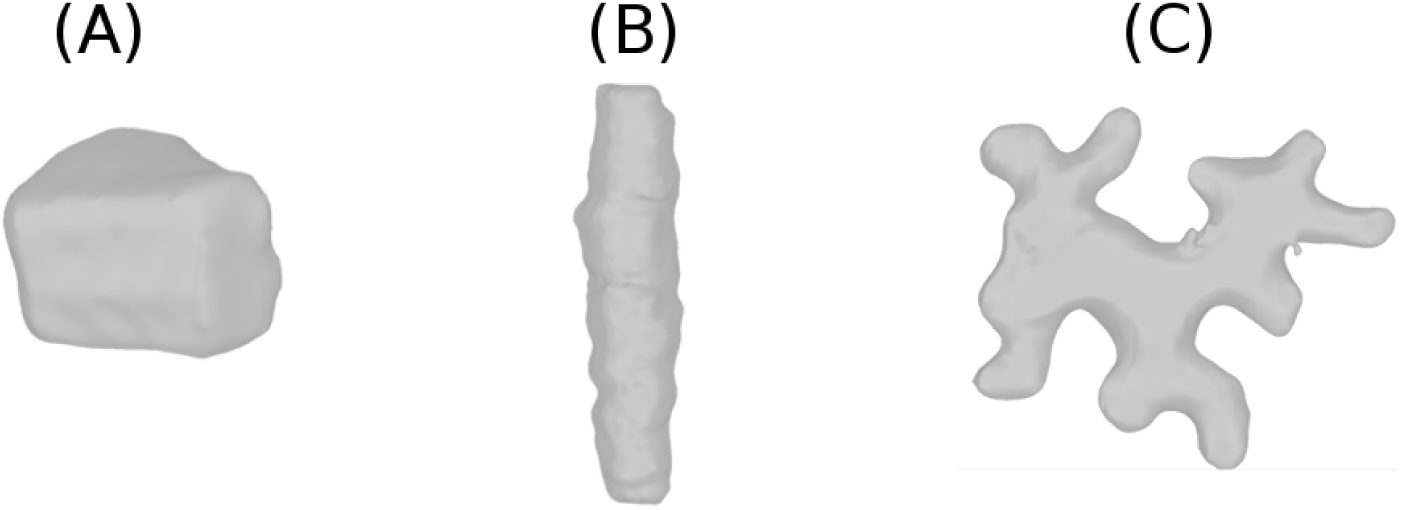
Plant cells with variety of sizes and shapes: (A) Arabidopsis root epidermal cell, (B) Sorghum leaf cell and (C) Tobacco leaf cell.

### Cortical MT dynamics

MTs are highly dynamic and filamentous protein polymer aggregates, and form one of the principal components of the plant cytoskeleton^11^. MTs have structurally two distinct ends - a minus end and a plus end. The plus end can dynamically switch from a growing state to a shrinking state or vice-versa. Switching of a MT plus end from a growing state to a shrinking state is called catastrophe while the reverse switching of a shrinking state to a growing state is called rescue. This phenomenon of reversible switching of MT plus ends between two states is called dynamic instability. On average, the minus end of an unstabilized MT continually is in a shrinking state. Thus, the combination of overall growth at the plus end and shrinkage at the minus end seemingly moves a MT as a whole. This motion is called treadmilling and has been observed in both in vitro^12–14^ and in vivo^15^. In contrast to animal cells, plant cells do not have a well defined MT organizing center. Instead MT activity is dispersed over the whole cell cortex, driven by the localized nucleation of new MTs by γ-tubulin complexes^16^. The cortical MTs are confined to a thin layer of cytoplasm just inside the plasma membrane of the plant cell and are attached to the cell envelope, ensuring that MTs do not translate or rotate as a whole^17,18^. In spite of their fixed attachment to the cell cortex, cortical MTs do show mobility which is due to treadmilling motion^17,19^. Twodimensional attachment to the cell cortex allows MTs to interact with each other via collisions, which occur when the polymerizing plus-end of a growing MT encounters a pre-existing MT. Depending on the value of the collision angle, three different possible events are observed^20^. For shallow angles (≲ 40^0^), a growing MT bends toward the direction of the MT encountered and this kind of adaptive event is called *zippering.* For steeper angles (≳ 40^0^), the encounter may lead to a so-called *induced catastrophe*, where the initially growing MT switches to a shrinking state. Alternatively, the growing MT may slip over the one encountered, leading to a *crossover* event.

In vivo imaging of cortical MTs has revealed that they nucleate at the cortex, either from isolated nucleation complexes or from pre-existing MTs, and gradually develop from an initially sparse and disorganized state into a final ordered array over a time period of an hour^17,18,21–24^ after the previous cytsokinesis. MTs have a finite lifetime and ultimately *disappear* by shrinking to zero length.

Plant cells within tissues typically have well defined relatively flat faces, bordered by edges of significantly higher curvature ^25^. In root epithelial cells microtubules have been observed to undergo catastrophes when they encounter these edges with a probability that increases with increasing curvature of the edge^26^. We will denote these as *edge*-*catastrophe* events (See Fig 2).

**Figure 2.**
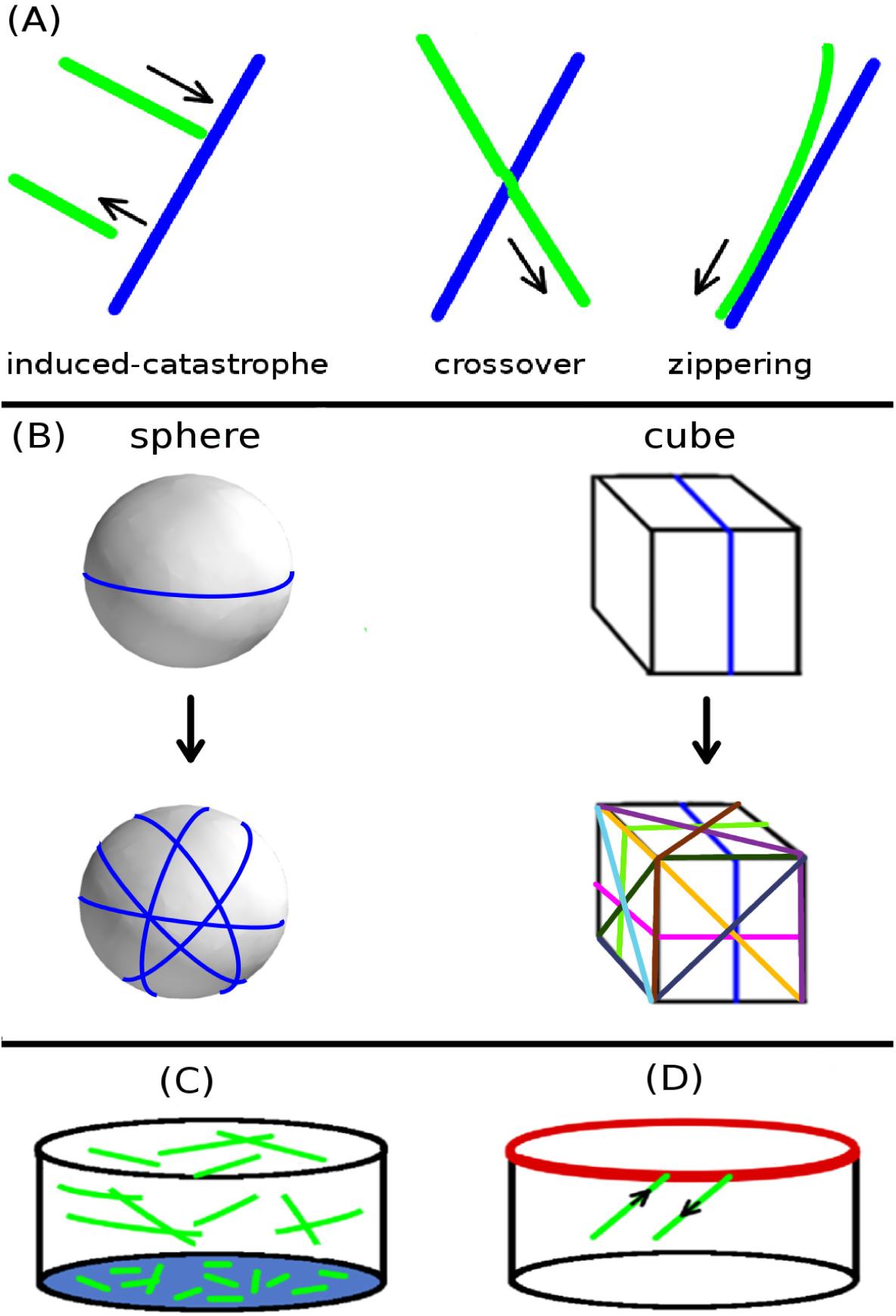
Different regulators of MT array formation, (A) Collision events during MT-MT interactions, where a steep angle of collision leads to induced-catastrophe or crossover, and shallow angle of collision leads to zippering.(B) Due to two dimensional attachment of the MTs to the cell surface (cortex), cell shape also influences MT interactions, hence MT order and array formation. On a spherical surface, an infinite number of closed geodetic paths are possible, suggesting an infinite number of possible paths of MT self-correlation. For a cubic surface, only nine such closed geodesics can be drawn, suggesting a limited set of possible paths of MT self-correlation. (C) Different degree of MT stabilization on different cell faces may lead to variation in the average length distribution of MTs, i.e. domain with enhanced MT stabilization will have longer average MT length than the domain of less MT stabilization (coloured in faint blue). (D) At an edge between cell faces (coloured in thick red), MTs undergo edge-catastrophe in a probabilistic manner, depending on the value of edge-angle.

### Models of CA formation

While existing experimental studies inform us about the molecular events and key parameters that are involved in CA formation^17,19,24,27,28^, it represents a complex emergent phenomenon that is the macroscopic result of a large number of stochastic microscopic events involving large numbers of MTs. For a full mechanistic understanding of biological processes of this type, mathematical and computational modelling has by now been recognised as indispensable^29^. From the outset, starting with the seminal work of Dixit & Cyr^20^, attempts have been made to model CA formation (for reviews on the different approaches involved please consult^30,31^). As far as the influence of cell geometry on CA formation is concerned, modelling studies have so far been limited to 2D approaches where the effect of a closed shape in 3D was mimicked by boundary conditions^32^, or geometrically simple shapes where boundaries are readily described by closed form mathematical expressions, such as cubes^33^ and cylinders^34^. Also, some of the simulations reported relied on finite time-step brownian dynamics, at most using a Gillespie-like algorithm for the spontaneous changes in MT dynamical state. This severely limits the time-efficiency of these simulations, and makes the acquisition of results for wide ranges of parameters with sufficient statistics a very difficult task. Tindemans et al.^34^ therefore developed a fully event-driven algorithm, in which not only the spontaneous stochastic state changes but also the collisions between MTs are performed by directly sampling from a dynamic list of potential future collision events and their associated waiting times, which is constantly updated. This achieves a speed up of at least three orders of magnitude with respect to standard finite time step algorithms, and eliminates errors in the location of the collision points as well.

### Our approach

Here we show how the algorithm of^34^ can be implemented efficiently on arbitrary triangulated surfaces, such as those obtained by segmenting 3D reconstructed confocal microscopy images of cells. Strikingly, the additional computational overhead incurred by having to determine the correct passage from triangle to triangle in a manner consistent with the 3D path, is almost completely balanced by the speed-up of the collision predicting algorithm due to the strong localization of the search area. In this way the new implementation is almost as fast as the original implementation, and allows the rapid exploration of many different parameter settings and/or geometries each sampled independently many times to obtain statistically reliable averaging over the stochastic ensemble. In addition, the MT dynamical parameters can be chosen independently within each triangular domain and passage probabilities can be associated with edges between domains. This allows us to easily implement biologically relevant effects such as the observed edge-catastrophes at locations of strong curvature, or potential differences in local MT dynamics due to developmental distinctions between cell faces, e.g. between faces created by recent cell divisions and faces that have expanded by growth.

## Methods

Here we outline our computational method for simulating MT dynamics on surfaces of arbitrary shape. First, we describe the processing of confocal laser scanning microscopy images of plant cells, transforming them into a triangulated surface mesh that can be directly used as simulation input. Next, we describe the implementation of the simulations and the order parameter that we define to measure the degree of MT order and the associated array orientation. Finally, we discuss how the working set of parameters for our simulations was selected.

### Processing confocal images for simulation

In Fig 3, we illustrate the procedure of transforming experimentally obtained confocal image into simulation input. First, using the image processing software *morphographX*^35^, we segment the experimentally obtained confocal images and extract the different cell shapes. Then, we approximate the cell shapes with an appropriate triangulated surface mesh. Using the surface mesh processing software *meshLab*^36^, we tag the different cell-edges and cell-faces, which are represented by appropriately coloured associated triangles. Finally, all this information is stored in a single mesh file, where each triangle has a cell edge-tag with the values of the associated cell edge-angles and cell face-tags (see Supporting Information Sec. SI.1)

**Figure 3.**
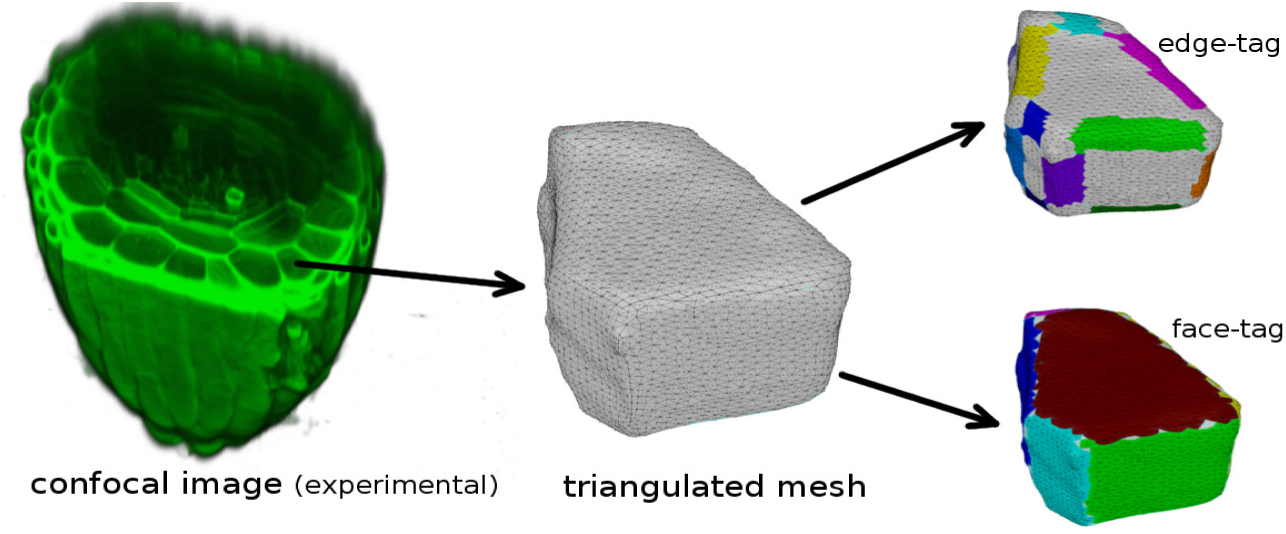
Image processing steps in transforming confocal image of a cell into a triangulated surface mesh, used as input to the simulation. Confocal images of the root apex of *Arabidopsis Thaliana* courtesy Viola Willemsen (WUR).

### Implementation of simulations

Confinement of the dynamics of cortical MTs to the surface (cortex) of plant cell effectively reduces the system to two spatial dimensions. In our modelling framework, we exploit this advantage by treating the cell shape as a two dimensional surface, embedded in the ambient there dimensional space. Because we are working with a triangulated approximation of surfaces, we developed an algorithm to establish a connectivity graph of the triangles^37^ in order to appropriately propagate MT dynamics between adjacent triangles. Using a combination of rotations, implemented by quaternions^38^, and *z*-axis translation, we transform the 3D-triangulated surface from the three dimensional *x* – *y* – *z* space to a set of disjoint 2D-triangles in the two dimensional *x* – *y* plane (see Fig 4.

**Figure 4.**
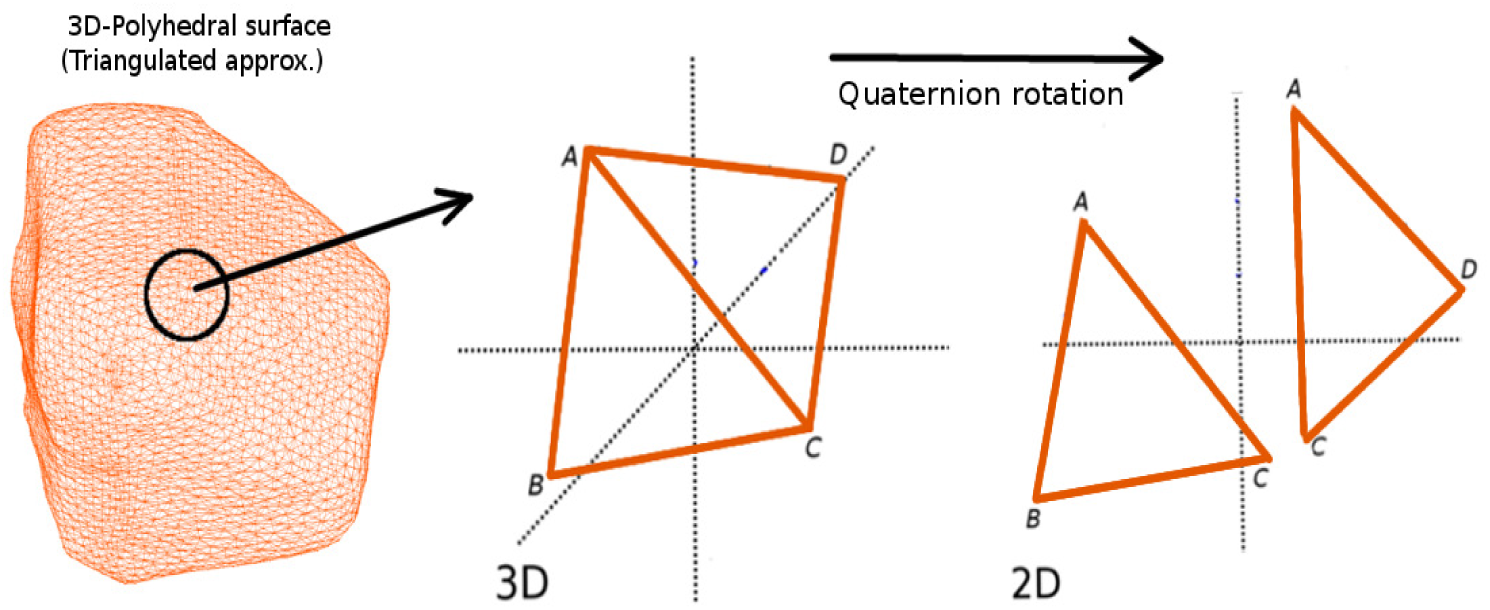
Mapping of triangles from the three dimensional surface mesh to the two dimensional plane. Δ*ABC* and Δ*ADC* are connected through a shared edge *AC* in three dimensional *x* – *y* – *z* space, but end up separated in the two dimensional *x* – *y* plane.

We simulate MT nucleation, growth, shrinkage and collision events within each individual triangle. In order to nucleate new MTs on the surface, we first randomly select a triangle from the entire set and generate uniformly distributed points within the selected triangle^39^. The initial MT growth direction is made isotropic by choosing a random angle of nucleation in the range [0, 2π]. The persistence length of a MT is much bigger than the average length^40,41^, allowing us to model the individual MT as an elongating straight line segment. The location of the collision point between two MTs is then readily determined by calculating the intersection point of their respective trajectories^42,43^ (see Fig 5). To deal with the various MT events, we implement an event-driven simulation technique^34^. The events are separated into two categories: stochastic events and deterministic events. The stochastic events associated with MTs are independent and result from Poissonian random processes specified by the nucleation rate *r_n_*, the spontaneous catastrophe rate *r_c_* and the rescue rate *r_r_*. The deterministic events associated with the MTs include collisions, disappearance, intersection with triangle edges, and simulation control events (i.e extracting simulation output at fixed time intervals). In between events, the plus-ends of MTs either grow or shrink with velocities *v*_+_ and *v*_−_ respectively, while the minus-end retracts with the treadmilling speed *v_tm_*, so that all length changes between events are readily computed.

**Figure 5.**
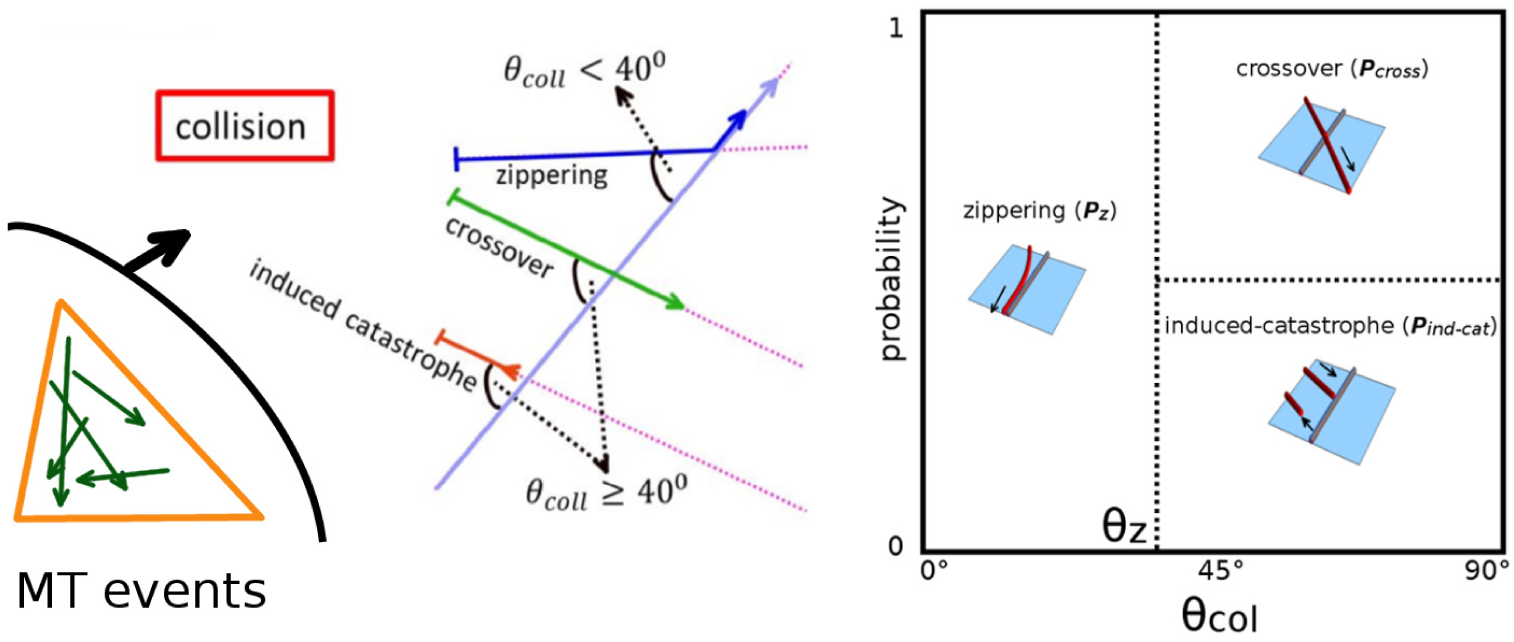
Simulation of MT events on an individual triangle. Upon nucleation, a straight line trajectory is assigned as a track for growth. Dynamic instabilities of MT tips are included through specification of spontaneous catastrophe and rescue rates. Type of collision events, i.e. zippering, induced-catastrophe and crossover, is determined on basis of the intersection angle between MT trajectories.

We use edge-to-edge links (obtained from the connectivity graph) to propagate MT dynamics from one triangle to its relevant neighbour. For example, in Fig 6 Δ*ABC* is connected with Δ*ADC* through the shared edge *AC*, which are linked. The proper transition of a MT from Δ*ADC* to Δ*ABC* through the edge *AC* on the three dimensional surface requires, in the two dimensional plane,

1. *Translation* of the associated growing MT tip from edge *AC*_(Δ*ADC*)_ of Δ*ADC* to edge *AC*_(Δ*ABC*)_ of Δ*ABC*.
2. *Rotation* of the MT trajectory from Δ*ADC* to Δ*ABC*.

Using an affine transformation^44^ on the coordinates of the growing MT plus end tip at point *P′* (*x′*, *y′*) of edge *AC*_(Δ*ADC*)_, we *translate* (Fig 6 left panel) the tip to the edge *AC*_(Δ*ABC*)_. Next, we *rotate* (Fig 6 right panel) the MT trajectory with rotation angle,

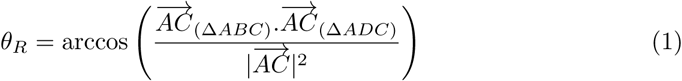

This rotation in the two dimensional plane is necessary to keep the MT growing along its original straight line path on the three dimensional surface. During transition of a MT from one triangle to another triangle, we implement the probability of local edge-catastrophe based on the local value of edge-angle between the adjacent triangles (see Supporting Information Sec. SI.2). We also use the face-tag, which is assigned to the triangles during the input surface mesh creation (see Supporting Information Sec. SI.3), to implement local differences in stability of MTs e.g. by changing the local catastrophe rate. More generically, our simulation framework allows arbitrary domain specific parametrisation, with the smallest domain being a single triangle. This flexibility allows full freedom to incorporate experimental observations on local MT dynamics or test additional relevant hypotheses.

**Figure 6.**
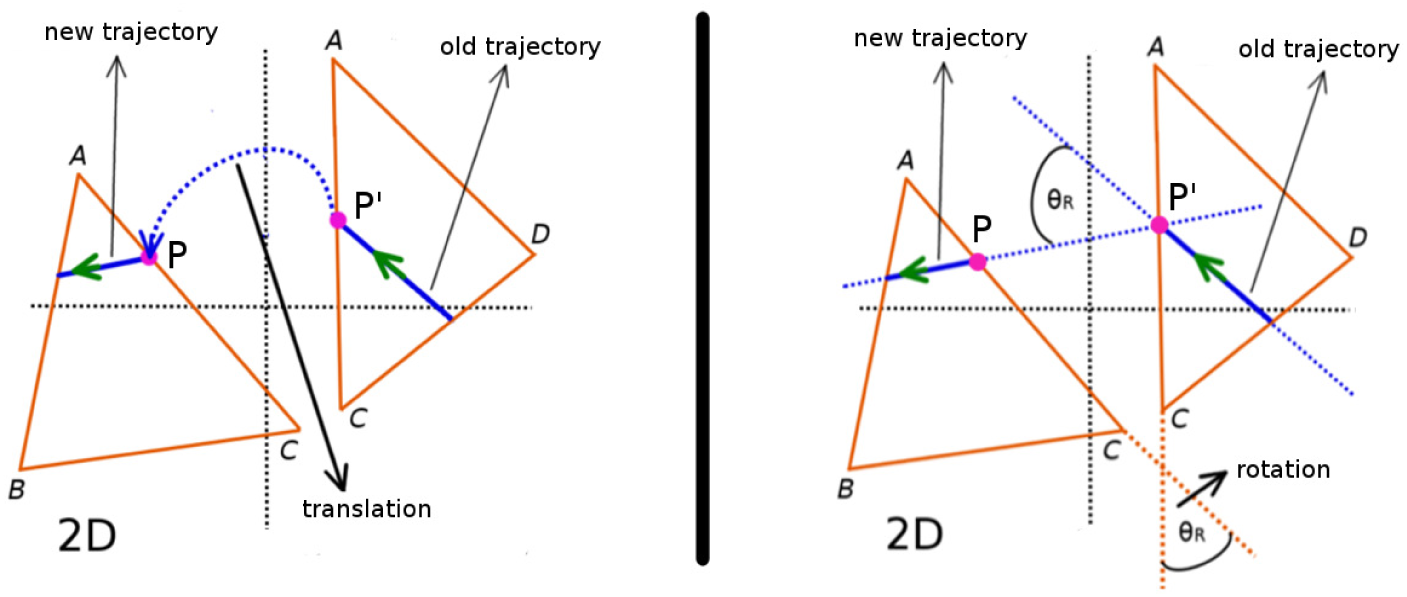
Propagating MT dynamics between neigboring triangles. An affine transformation is used to translate a growing MT tip from edge *AC*_(Δ*ADC*)_ of Δ*ADC* to edge *AC*_(Δ*ABC*)_ of Δ*ABC* (left panel). Upon translation, the MT growth direction is rotated by an angle *θ_R_* (see Eq 1) that equals the angle between *AC*_(Δ*ADC*)_ and *AC*_(Δ*ABC*)_. A new trajectory along this rotated direction is then generated for the MT to continue its dynamics on Δ*ABC* (right panel).

Finally, whenever required we apply the inverse transformation on the set of two dimensional triangles and the MT segments present on them, to reconstruct the three dimensional surface with the MT configurations realised. Through this transformation, segments of a MT which appear to be disconnected in the set of two dimensional triangles, get (re)connected on the three dimensional surface. As illustrated in Fig 7, the segment *M*_1_*M*_2_ is attached to Δ*ABC* and the segment *M*_2_*M*_3_ is attached to Δ*ADC* in two dimensional plane. After the inverse mapping, these two segments get connected in the three dimensional surface and represent a single MT with end points (*M*_1_, *M*_3_).

**Figure 7.**
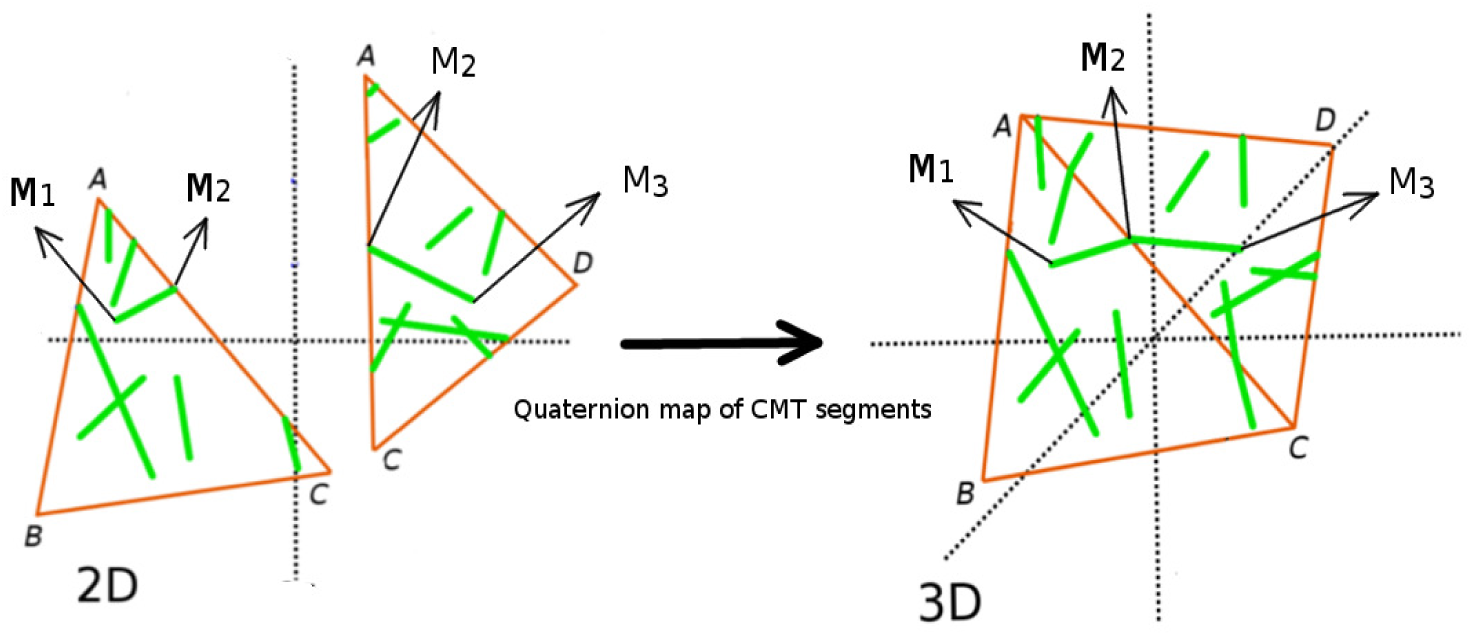
Mapping MT segments from two dimensional plane back to the three dimensional surface. The inverse quaternion rotation operator of Δ*ABC* is applied to the segment *M*_1_*M*_2_, and a similar operation is performed on MT segment *M*_2_*M*_3_ of Δ*ADC*. If Δ*ABC* and Δ*ADC* are connected to each other via the edge *AC*, the MT segments *M*_1_*M*_2_ and *M*_2_*M*_3_ also get reconnected in their three dimensional position.

To measure the degree of order and the orientation of the MT array, we define two order parameters. The first parameter is a scalar (*Q*^(2)^), which measures the average
*degree* of ordering of the MTs, and the second a vector Ω̂, which indicates the global *orientation* of the array. Both these parameters are derived from a single tensorial order parameter **Q**^(2)^ (see Supporting Information Sec. SI.4), *Q*^(2)^ being the absolute value of the smallest eigenvalue of this tensor and Ω̂ the corresponding normalized eigenvector, i.e. |Ω̂| = 1. For completely random orientation of MTs *Q*^(2)^ ≈ 0 and for well organised MTs that form an array, *Q*^(2)^ ⪅ 1. The direction of Ω̂ is *perpendicular* to the average local orientation of the MTs (see Fig 8), i.e. perpendicular to the plane onto which the total projection of individual MTs is maximal on average.

**Figure 8.**
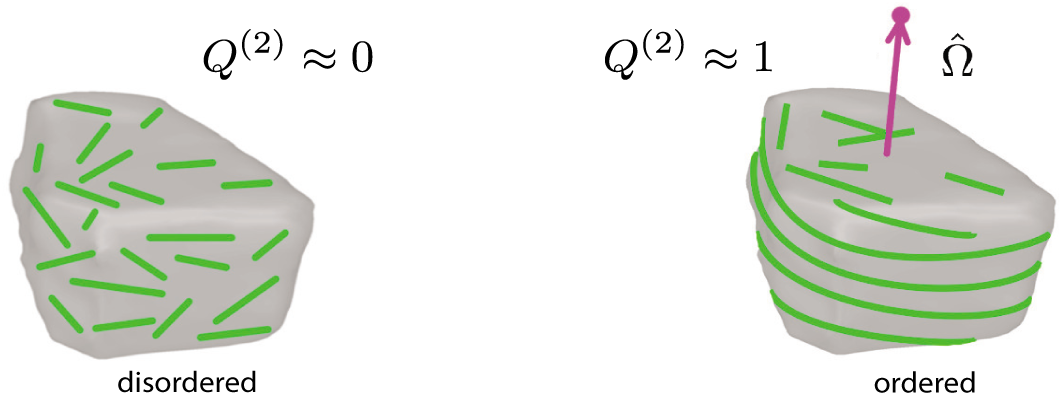
Schematic representation of the global order parameter. The scalar part *Q*^(2)^ measures the degree of MT order. The vector part Ω̂ measures the orientation of the associated MT array and is perpendicular to the plane of the MT array (magenta arrow). The dot (a filled magenta circle) at the tip of Ω̂ provides a concise representation of the MT array orientation.

In Fig 9, we present an overview of the entire simulation approach.

**Figure 9.**
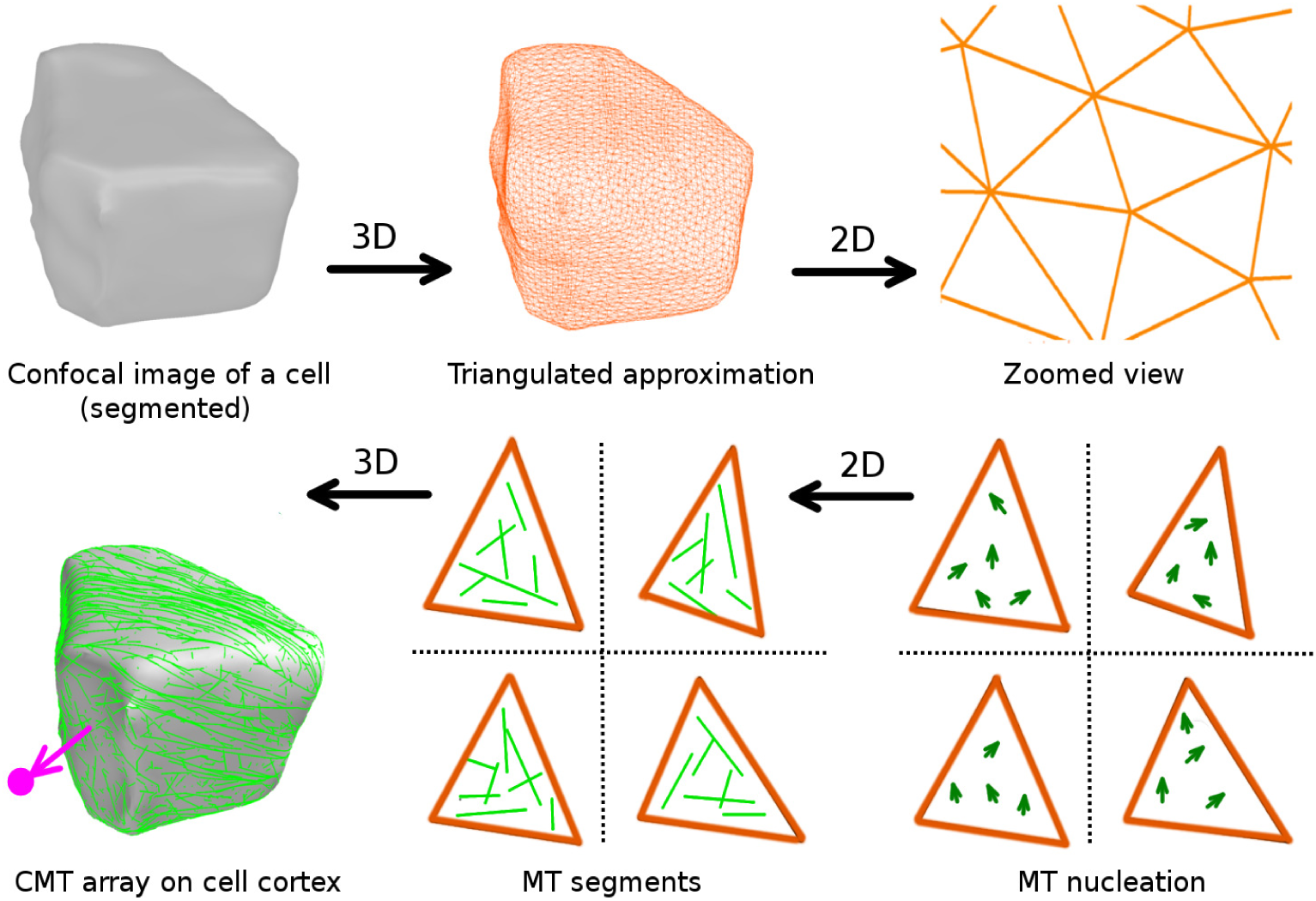
A schematic overview of the workflow in our simulation approach.

### Working domain of parameter values for simulation

In order to obtain a suitable set of MT dynamical parameters for our simulations, we make use of the control parameter *G* introduced and used in^45,46^. This parameter encapsulates the effect of all six single MT dynamics parameters (speeds *v*_+_,*v*_−_ and *v_tm_*, dynamical instability rates *r_c_*,*r_r_* and nucleation rate *r_n_*) into a single magnitude that controls the frequency of MT interactions which are required to ensure a spontaneously ordered state. Its explicit form identifies it as the ratio between two length scales

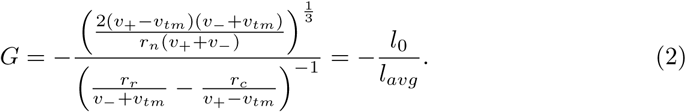

Here *l*_0_ can be interpreted as the typical distance between MT encounters, and *l_avg_* is the mean length of MTs in the absence of any interaction effects. The statement that *G* is the single control parameter was only strictly derived for spatially homogeneous and unbounded domains. To allow for possible effects of a bounded cell geometry, we chose to independently vary the two factors involved: *l*_0_ by varying the MT nucleation rate *r_n_* and *l_avg_* through the spontaneous catastrophe rate *r_c_*. Moreover, and unless explicitly mentioned, we implement homeostatic control of the MT growth speed *v*_+_ by implementing a finite tubulin pool (see Supporting Information Sec. SI.5 for details), as this speeds up the relaxation towards a steady state. The remaining simulation parameters are taken from^34^. We tested the parameter choice on a spherical cell with a radius *r* ≈ 6 *μm*, inspired by the typical dimensions of a *Arabidopsis thaliana* embryonic cell. This is at the smaller end of the spectrum of plant cell sizes and therefore the most stringent test on finite size effects. We found that we could obtain robustly ordered arrays with *Q*^(2)^ ≳ 0.70 at a state point with *l*_0_ = 2.05 *μm* and *l_ave_* = 417.5 *μm*, corresponding to *G* ≈ −0.005 (considering no MT-MT interaction, i.e. prior simulation in Fig 10). Variation in the value of average MT length in the presence of MT-MT interaction (i.e, average MT length measured in simulation and denoted by 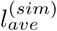) with respect to *l_avg_* is described in Supporting Information Sec. SI.5. The fact that *G* < 0 ensures that we are in the bounded growth regime, were the length of MTs is always finite. All parameters used are summarised in Table 1 (see Supporting Information Sec. SI.8).

**Figure 10.**
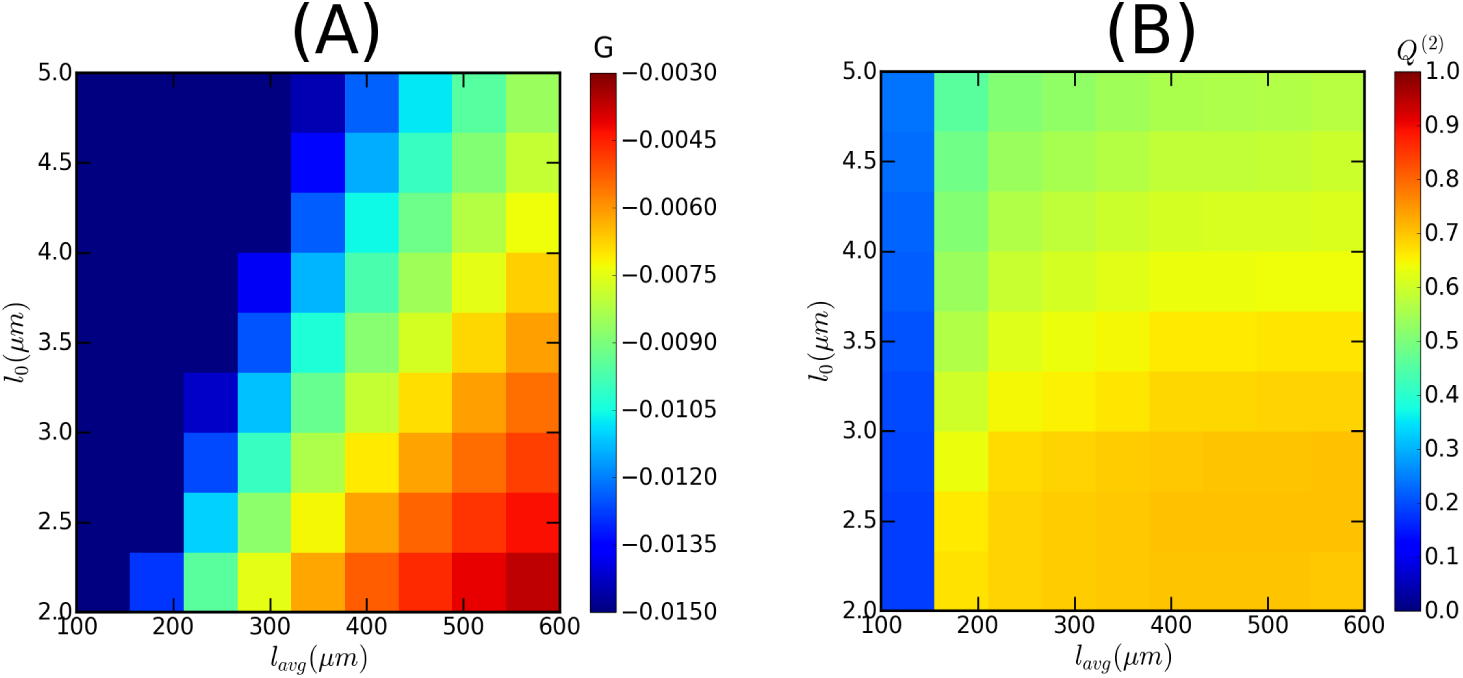
Working domain of MT simulation. Heat map showing: (A) The values of MT-MT interaction control parameter *G*, and (B) The values of the global order parameter *Q*^(2)^. A value of *G* ≲ −0.005 assures sufficient MT-MT interaction to achieve an order parameter value *Q*^(2)^ ≳ 0.70 (for the time evolution, refer to Fig SI.5). The values of *G* were calculated by using the parameter values described in^46^, except the nucleation rate (*r_n_*). To assure sufficient number of MTs, we choose *r_n_* ≈ 0.01 *sec*−1 *μm*^‒2^. These simulations were performed using finite tubulin pool, *ρ_tub_* = 10 *μm*^μ1^.

## Results

### Validation of the framework

We performed a set of simulations validating our computational framework. We first tested to what extent the triangulation of otherwise smooth surfaces influences the final results. To do so we considered the case of approximating a perfect sphere, varying both the number of triangles employed, as well as the triangulation method. The key desiderata in this case are (i) that in steady state, due to the spherical symmetry, there should be no bias in the orientation Ω̂ of the ordered array, and (ii) that the value of the scalar order parameter *Q*^(2)^ does not depend on the nature of the triangulation. Next, in order to test the system as a whole against a known, and geometrically non-trivial case, we reproduced previously reported results on a cubic surface^47^.

### Effect of triangulation

We triangulated a spherical surface with numbers of triangles ranging from *T* =10 − 5000, using four different triangulation algorithms. On each of these geometries we ran 1000 independent realizations of the stochastic simulation. Generically, the steady state achieved in these simulations is a diffuse equatorial band of ordered MTs. To test for isotropicity, we considered the distribution of the normalized vectorial order parameter Ω̂. This showed that for triangle number *T* > 100 the isotropicity is already satisfactory. We illustrate this in Fig 11 (A-C) with the results for *T* = 5000, which is the order of magnitude of the number of triangles obtained by the meshing of the confocal microscope images. Looking at the steady values of the scalar order parameter *Q*^(2)^, we found that, except for the smallest number of triangles *T* = 10, the results for the time to reach steady state as well as the steady values were largely independent on the triangle number, as illustrated in Fig 11 (D). Moreover, these results did not depend on the specific triangulation algorithm used (For details please refer to the Supporting Information Sec. SI.7).

**Figure 11.**
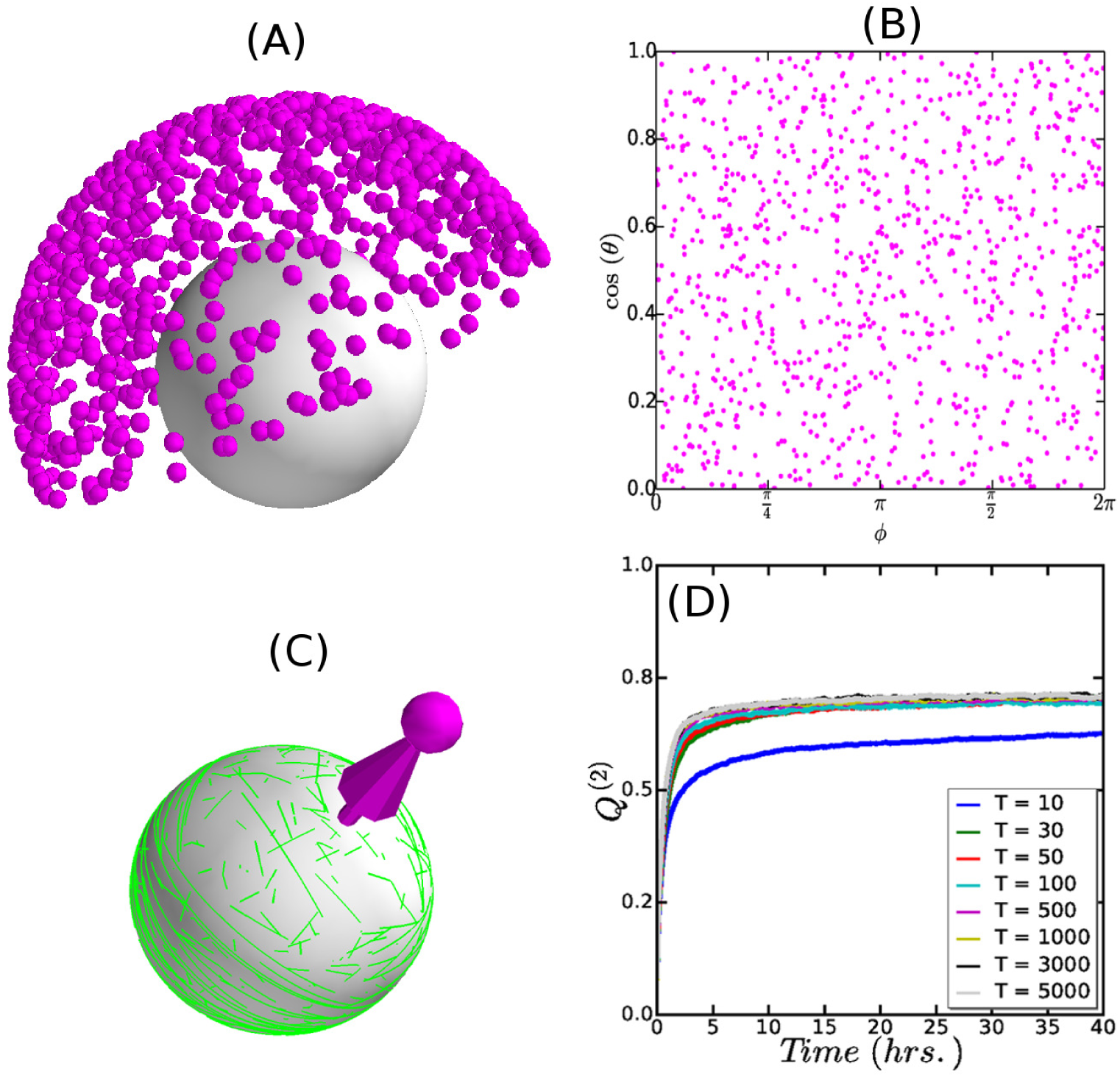
(A) Simulated orientation of MT array on a spherical surface. Simulations were performed for ≈ 1000 independent realizations of the stochastic MT dynamics. Each dot point represents the tip of the orientation vector Ω̂ associated with a simulation configuration, and perpendicular to the associated MT array. (B) Mapping the angular components (*θ*, *ϕ*) of the spherical coordinates of the Ω̂ tips to the equivalent two dimensional plane (*x*,*y*) (see Supporting Information EqSI.18). (C) A simulation snapshot of the MT array associated to the specific Ω̂ tip. (D) A comparison in the time evolution of the degree of MT order (*Q*^(2)^) for the different triangulations, reflecting robust reproducibility in the degree of MT order for *T* > 30.

### Simulation on cubic surface

To test the ability of our simulation framework to reproduce existing MT simulation results, we considered a cubical geometry. In the original simulation reported in^47^, the cube was built up out of 6 square planar domains. Here, we triangulated the cube by subdividing each face along the diagonals into 4 isosceles triangles. The edges between the faces were classified into two groups: the ones bordering the top and bottom faces are denoted, paraphrasing the standard nomenclature for plant root cells, as periclinal, shown in Fig 12 panel A edges with solid lines, while the remaining, dotted, edges are called anticlinal. Edge-catastrophe probabilities are associated to the edges, denoted by *P_PC_* and *P_AC_* for the periclinal and anticlinal edges respectively. Following^47^, we adopt the following orientational order parameter 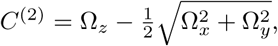 where the *z*-axis is chosen perpendicular to the top and bottom face. A value of *C*^(2)^ = 1 corresponds to prefect anticlinal orientation of the array, while *C*^(2)^ = −1/2 signals a periclinal orientation. The intermediate value *C*^(2)^ = 0 corresponds to a mixed state with an equal probability for each of the three principal ordering directions. To obtain the unbiased state, we first considered *P_PC_* = *P_AC_* = 0.26, the value reported in^33^ for the catastrophe probability at the periclinal edges in *A. thaliana* root epidermis cells. As Fig 12 panel B shows, we find a distribution of *C*^(2)^ values, with roughly 2/3 with a value of *C*^(2)^ ≈ −0.5 corresponding to the two equivalent periclinal orientations, and 1/3 with *C*^(2)^ ≈ 1, corresponding to the anticlinal orientation. We then keep *P_PC_* = 0.26 fixed, and scan *P_AC_* in the range 0 − 1 from high to low values. We see (Fig 13) that initially the system remains locked into the anticlinal orientation characterized by *C* ≈ 1. When the value of *P_AC_* become comparable to *P_PC_* (0.2 ⪅ *P_AC_* ⪅ 0.3) the system enters a transition regime in which it can stabilize to either the anticlinal or the periclinal state with relative frequencies dependent on the precise value of *P_AC_*. Finally, for *P_AC_* ⪅ 0.1, only the periclinal orientation (with equal probability in both of the possible directions), was observed, i.e. a state flipped by a complete 90° degree flip from the original anticlinal orientation. The results are fully consistent with those reported previously^47^, including the occurrence of an additional set of diagonal MT orientations when edge-catastrophes are absent (see Supporting Information Sec. SI.6).

**Figure 12.**
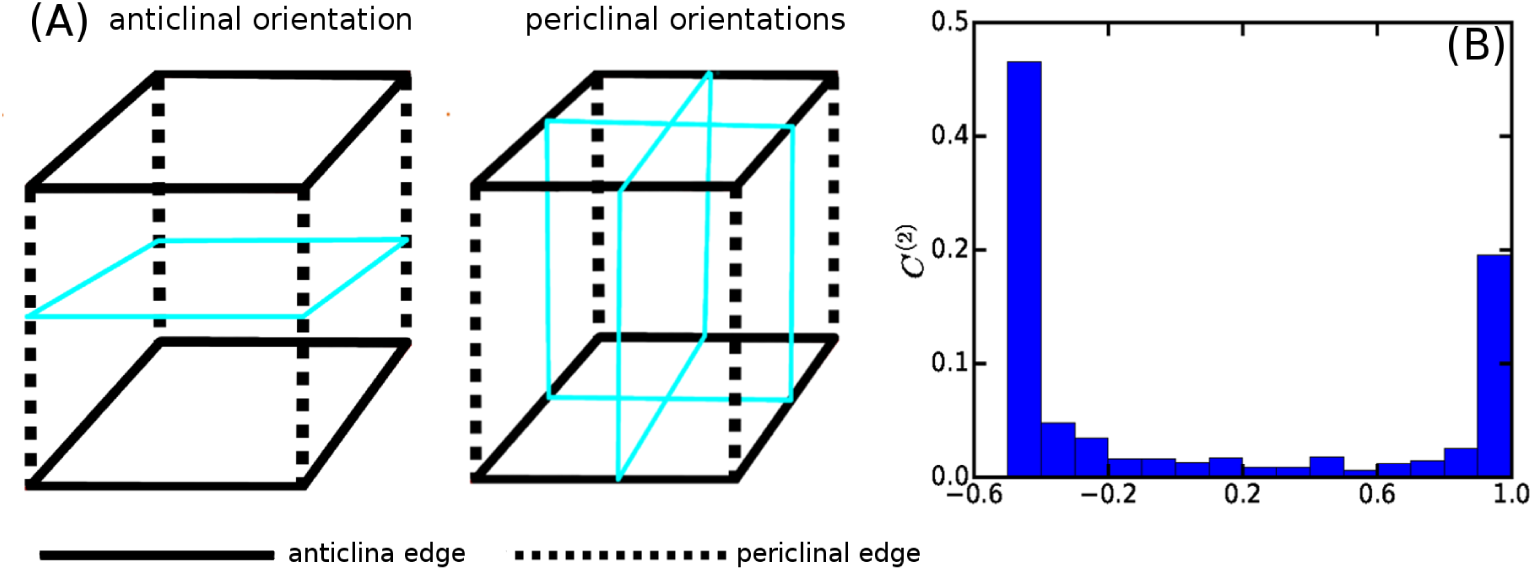
(A) Schematic diagram of anticlinal (horizontal plane in cyan color) and periclinal (vertical planes in cyan color) orientation direction and tagging the edges for the implementation of edge-catastrophe. The anticlinal edges are tagged by solid line and the periclinal edges are tagged by dotted lines. (B) Distribution of *C*^(2)^ values for *P_AC_* = *P_PC_* = 0.26, which is a bimodal distribution with peaks at *C*^(2)^ ≈ 1 and *C*^(2)^ ≈ −0.5, obtained from ≈ 1000 independent realizations. In these simulations we used an infinite tubulin pool and the same value of *G* as described in^47^.

**Figure 13.**
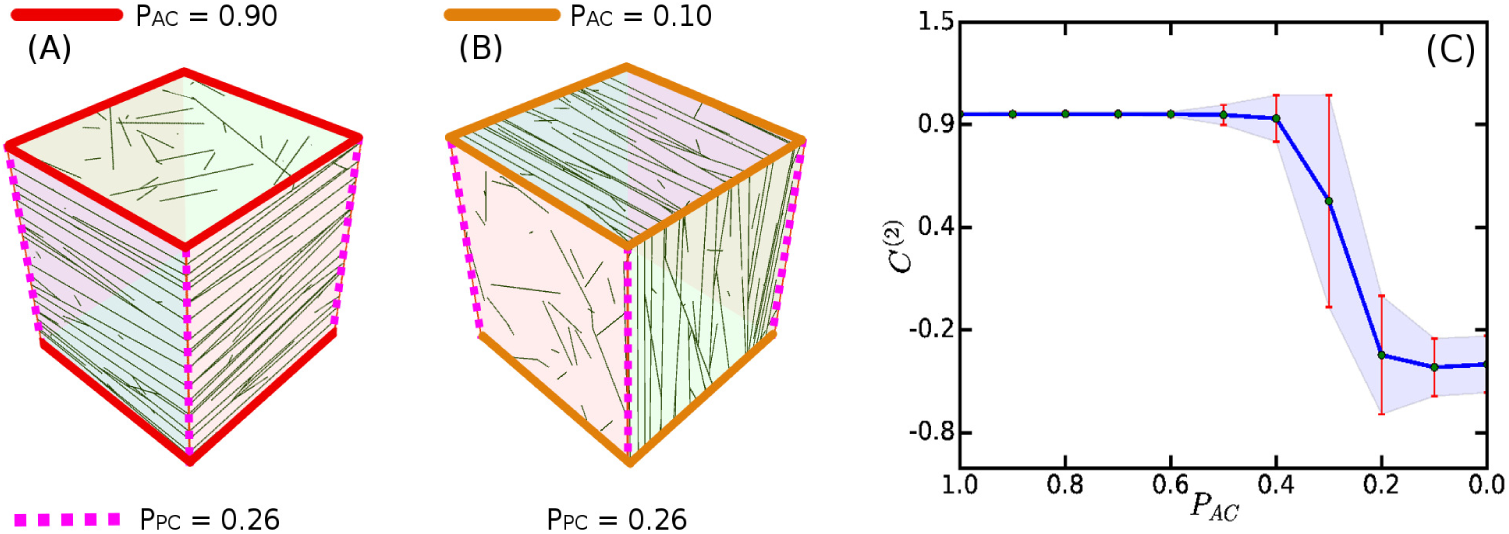
Simulation of MT dynamics for fixed value of *P_PC_* = 0.26. Simulated orientation of the MT array for: (A) *PA_C_* = 0.9 (anticlinal) and (B) *PA_C_* = 0.1 (periclinal). (C) Gradual increase in the probability of switching of the MT array orientation for anticlinal (*C*^(2)^ ≈ 1) to periclinal (*C*^(2)^ ≈ −0.5). Each point in the plot is an average of ≈ 1000 independent simulations. Parameter values as described in^47^.

## Probing the influence of cell shape on MT order

We are now in a position to use our framework to study the role of cell shape on array formation. We first do this in a geometrically simple setting, which allows us to systematically disentangle the role the various shape-related factors involved. Then we turn a real cell shape, provided by the leaf pavement cells of *Hedera helix.*

### Interplay between shape effects and stability rules

As the example of the cubical surface discussed above shows, shape by itself already has an impact on MT organisation, essentially selecting a finite number of possible stable array orientations. Adding edge-catastrophes can then serve to uniquely stabilize one of these possibilities. There is, however, a third factor that can play a role namely differences in MT dynamics localized to specific cell faces. The possibility of cell face-dependent MT dynamics has been discussed in the context of array organisation due to environmental cues such as light^48^ and hormone stimulation^49^. Such differences could potentially also derive from developmental difference between the cell, e.g. the difference between faces created by division versus expanding cell faces, or maturation effects due to “ageing” of the associated cell wall. Here we assess the interplay between all these factors in a non-trivial, yet simple cell shape. We triangulated a rectangular parallelepiped of dimensions *a* = 13 *μm*,*b* = 7.5 *μm* and *c* = 5 *μm*, subsequently rounding its edges using *meshLab*^36^. These dimensions, and the resultant cell volume, were chosen to provide a highly stylized version of an early stage A. *thaliana* embryonic cell. The MT dynamical parameters were chosen from the working domain of simulation to yield a steady state mean MT length with MT-MT interation to be 4.5 *μm*, ensuring that MTs are sensitive both to edge and shape effects. To discuss the possible orientations of the MT array, we introduce three unit vectors *p̂_a_*,*p̂_b_* and *p̂_c_* parallel to corresponding edges (see Fig 14(A)). In the default case, i.e. without any additional effects, we find that in 51% of all realizations the vector order parameter aligns along the *p̂_a_* direction, in 46% along the *p̂_b_* direction, and the remaining few percent in spurious directions, but notably none along the *p̂_c_* direction. This is readily understood as the array orientations in this finite geometry are stabilized by correlations due to avoidance of self-interactions. These correlations are stronger along shorter closed paths around the shape. This immediately predicts that a parallel MT array perpendicular to *p̂_a_*, which covers the shortest circumference *C_a_* = 2(*b* + *c*) = 12.5*μm*, should be most favoured, followed by the orientation *p̂_b_*, with *C_b_* = 2(*a* + *c*) = 18*μm*, whereas the least favored direction *p̂_c_* with *C_c_* = 2(*a* + *b*) = 20.5 *μm* apparently is no longer stabilized. Implementing edge-catastrophes as described in Supporting Information Sec. SI.2 with edge-catastrophe multiplier *E_cat_* = 2 to all edges completely reverses this tendency. Directions in which MTs would on average cross more catastrophe-inducing edge crossings over their innate length become unfavorable. In this case the orientation *p̂_c_* with mean distance between edge crossings *d_c_* = 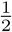(*a* + *b*) = 5.125*μm* is most likely (71%), followed by *p̂_b_* with *d_b_* = 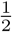(*a* + *c*) = 4.5*μm* at 28%, whereas now *p̂_a_* with *d_a_* = 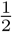(*b* + *c*) = 3.125*μm* does not occur at all. Finally, we considered the role of face stabilization. To that end we “protected” the faces with the surface area (*a* × *c*) and increased the MT catastrophe rate *r_c_* on all other faces such that the average innate length of the MTs on those faces decreases 3.5 *μm*. This obviously favors arrays which pass over the face with sides *a* and *c*. Indeed the results now show an uneven split between the orientations given by *p⃗_a_* (77%) and *p⃗_c_* (19%). In this case the direction *p⃗_b_* is explicitly suppressed as it would require MTs to pass through faces with lower stability.

**Figure 14.**
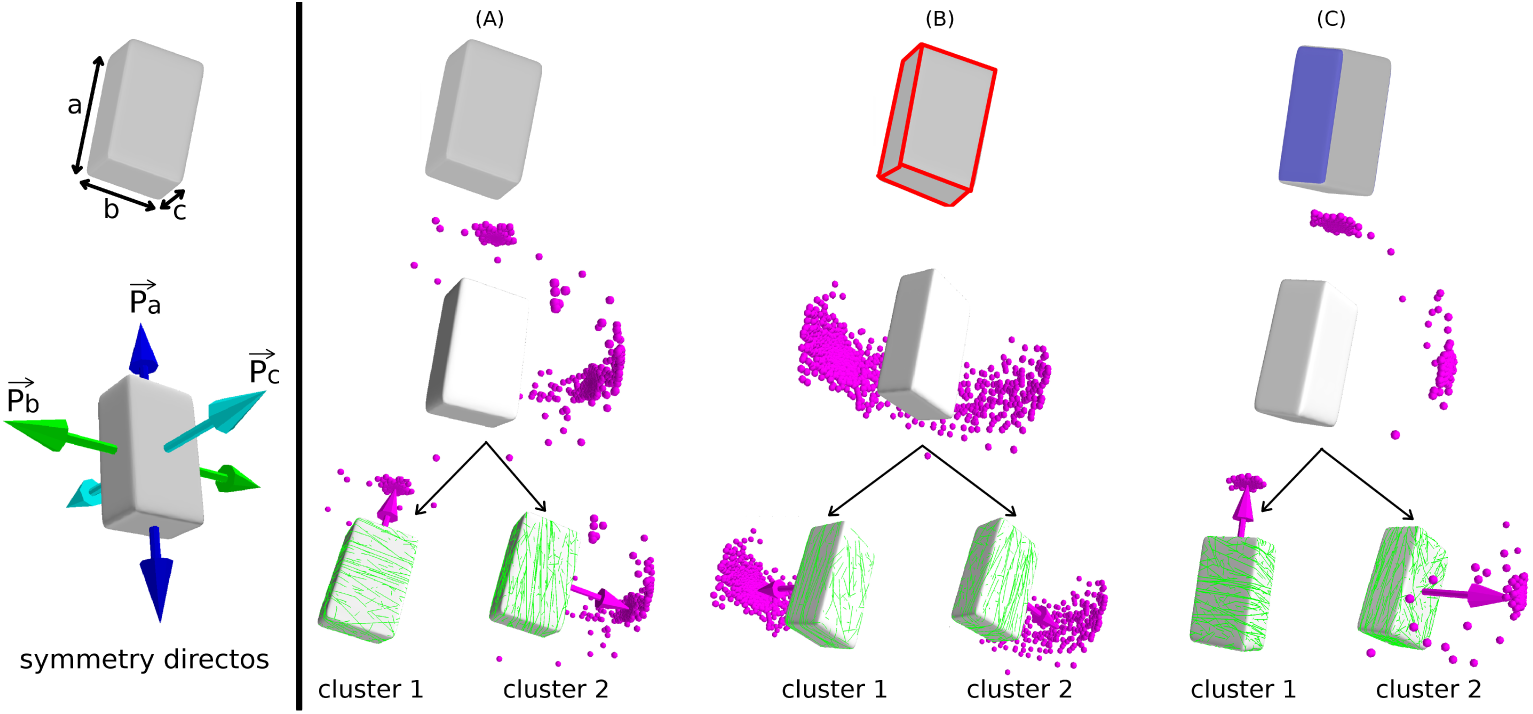
A schematic representation of a rectangular parallelepiped and the symmetry direction *p⃗_a_*,*p⃗_b_* and *p⃗_c_* parallel to the edges *a*, *b* and *c* respectively (left panel). Simulated orientation of MT arrays on the surface of the rectangular parallelepiped, where each dot point represents the tip of an orientation vector associated with a simulation configuration, and perpendicular to an associated MT array (right panel). MTs were simulated by considering three different regulatory effects in the dynamics of MT: (A) default shape, which resulted in two clusters (cluster 1: ≈ 52% [519/991] around *p⃗_a_*, cluster 2: ≈ 46% [452/991] around *p⃗_b_*) of MT array orientation, (B) edge-catastrophe, which resulted in two clusters (cluster 1: ≈ 28% [276/992] around *p⃗_b_*, cluster 2: ≈ 71% [706/992] around *p⃗_c_*), and (C) enhanced MT stabilization at a selected face, which resulted in two different clusters (cluster 1: ≈ 77% [762/995] around *p⃗_a_*, cluster 2: ≈ 19% [186/995] around *p⃗_b_*) of array orientation. For each of three cases of the three regulatory effects, simulations were performed for ≈ 1000 independent realizations.

Taken together these results show that both edge-catastrophe and face stability can independently exert a degree of control over the global array state.

### MT simulation on realistic plant cell shape

Although of varying degree of regularity (cf. Fig 1), some plant cells have highly nontrivial shapes. A case in point are the shapes of the leaf pavement cells of flowering plants such as *Nicotiana benthamiana* and *Hedera helix*, which have multiple highly irregular protrusions. As such they provide an ideal test case for our framework. We first segmented confocal images of such cells, produced a triangulated approximation to their shapes, followed by manual tagging of edges and faces when applicable. First, we simulated the MT dynamics on the resultant triangulated networks of the leaf pavement cells of *Nicotiana benthamiana* under default conditions. This resulted into almost random distributions of the global ordering orientation, restricted, however, to the semi-2d plane of the shape itself. Snapshots reveal a significant degree of local order, also reflected in the value of the global scalar order parameter *Q*^(2)^ ≈ 0.45. Local order included in several cases band formation around the necks of protrusions (see Fig 15 A). Such ringlike MT structures have in fact already been reported in *A. thaliana*^50^, where evidence was presented that actin is involved in localising these structures. The requirement for actin to constrain MT bands around the necks is consistent with our result that MT dynamics influenced by geometry alone cannot exclusively generate these “neck” configurations. As the snapshots in 15 panel A show, stochastic differences between individual realisations of steady state configurations lead to varying distributions of banded structures over the shape, which also explains the essentially random global array orientations in our simulations.

**Figure 15.**
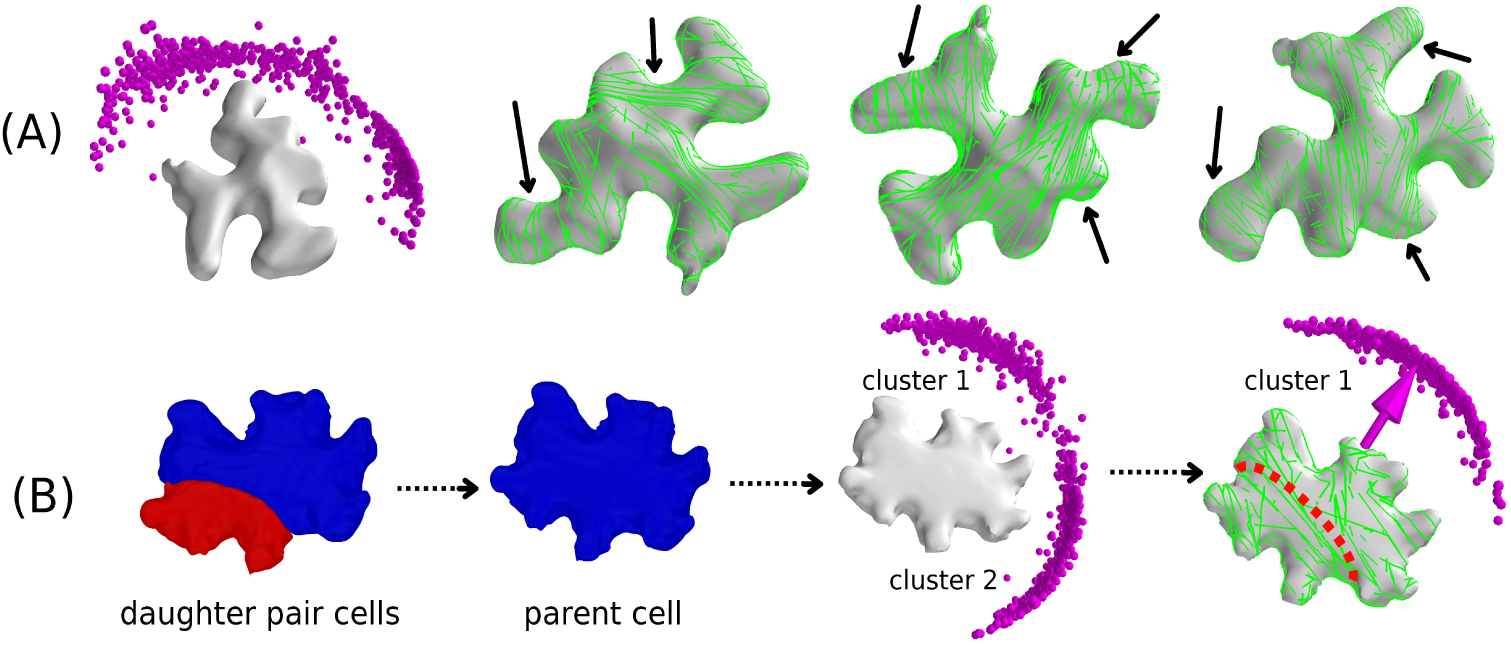
Simulated orientation of MT arrays on the surface of (A) *Nicotiana benthamiana* and (B) *Hedera helix* leaf pavement cell shapes, where each dot point represents the tip of an orientation vector associated with a simulation configuration, and perpendicular to an associated MT array. Simulations were performed by considering basic MT-MT interactions on default cell shapes: (A) Simulated orientation of MT-arrays on the leaf pavement cell of *Nicotiana benthamiana.* Resulting orientation is almost a random distribution, which is restricted on the semi-2d plane of the cell shape and reflects formation of local MT-arrays around the various irregular protrusions (Three different instances are shown). (B) A parent leaf pavement cell shape of *Hedera helix* is recreated by merging the daughter pair of cells, then MT dynamics was simulated on the default shape which resulted in two clusters of MT-array orientation with almost identical probabilities (cluster 1: ≈ 51% [509/991] and cluster 2: ≈ 49% [482/991]). Average MT-array orientation associated with the cluster 1 matched with the corresponding orientation of the division plane that made the daughter pair of cells (rightmost image).

In addition, we can now look for a possible correlation between a robust prediction of global orientation of the MT-array and the subsequent orientation of division plane^51,52^, by simulating MT dynamics on the triangulated network of a reconstructed parental *Hedera helix* leaf pavement cell. We used *morphographX*^35^ to recreate such a “parent cell”, by merging the corresponding daughter cell pair. The simulations revealed two distinct possibilities for average global ordering orientation as shown in panel B of Fig 16, with very similar frequency of occurrence: cluster 1 ≈ 51% and cluster 2 ≈ 49%. cluster 1 is slightly more compact, indicating a smaller standard deviation in the corresponding orientation. Strikingly, the orientation of cluster 1 corresponds to the actual orientation of the division plane that generated the cell pair. This shows that, at the very least, cell geometry alone, in the absence of additional controlling mechanisms, can predispose array orientation towards selecting division planes. Note that our simulations also appear to rule out the possibility of out-of-plane array orientation, consistent with the fact that periclinal divisions (parallel to the leaf surface) are extremely rare in leaf epidermis cells (maize ^53^, tobacco ^54^).

**Figure 16.**
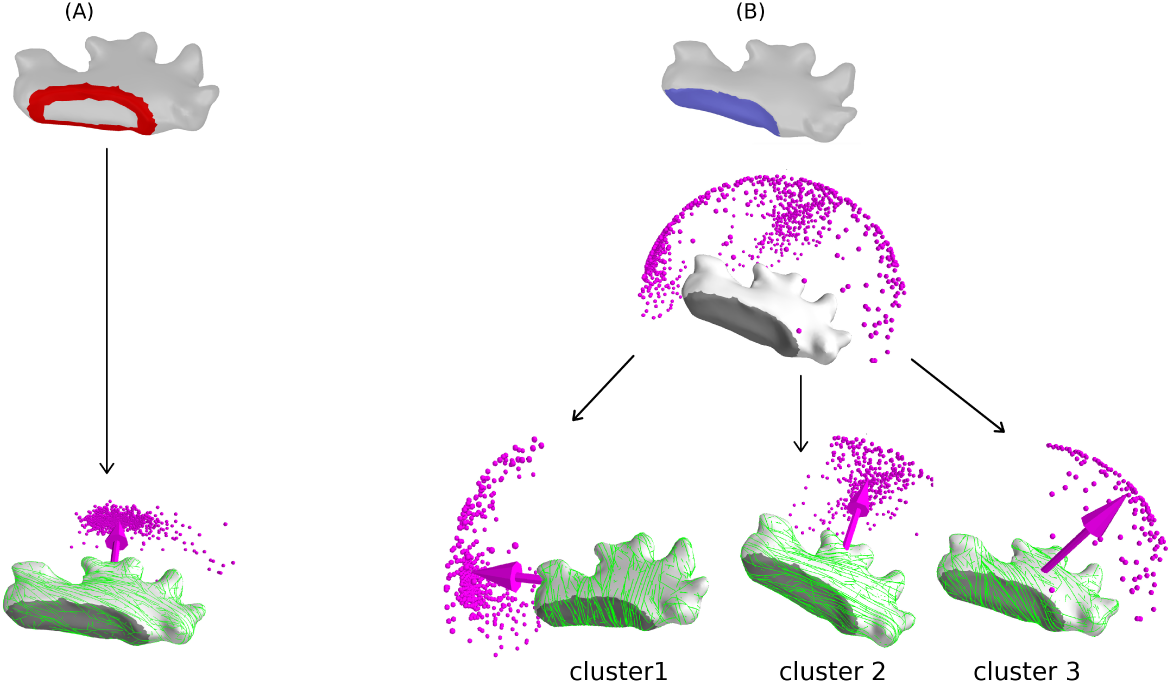
Simulated orientation of MT arrays on the surface of *Hedera helix* leaf pavement cell shape, where each dot point represents the tip of an orientation vector associated with a simulation configuration, and perpendicular to an associated MT array (right panel): (A) Simulation with edge-catastrophe, which resulted in only a single cluster of MT array orientation. Edge of the cell is coloured red to indicate that in simulation, the MTs were subjected to edge-catastrophes, and (B) Simulation with enhanced MT stabilization at the newly inserted cell face, which resulted in the formation three clusters of MT array orientation. Majority of the orientations were distributed in two nearly identical clusters (cluster 1: ≈ 37% [370/993] and cluster 2: ≈ 49% [484/993]). A small fraction of the MT array orientation formed a third cluster (cluster 3: ≈ 14% [139/993]), which depicts the effect of enhanced MT stabilization. For each of three cases of the three regulatory effects, simulations were performed for ≈ 1000 different configurations. Generally, these cells are bigger than the embryonic cells, however we rescaled their volume to the average volume of the embryonic cells, without affecting their shape. This allows us to use simulation parameters from the previously defined working domain, ensuring formation of ordered MT-array in the same time scale. Cell templates used in these simulations were provided by Ikram Blilou (WUR).

Finally, we applied our simulation frame work on a most recent generation of *Hedera helix* pavement cell, i.e a daughter cell of a just divided cell. These cells contain an edge and two developmentally distinct cell faces, thus providing a suitable realistic sample to explore the possible consequences of edge-catastrophe and MT-stabilization on the MT-array orientation. Simulation with edge-catastrophes to the circumference of the basal cell face uniquely selects only a single average orientation of MT-array (see panel B in Fig 16). This again highlights the ability of cell intrinsic features, such as the sharp edges created by a preceding division plane, to uniquely determine array orientation, and therefore potentially the subsequent division plane. However, it is conceivable that for developmental purposes the cell would require additional control, beyond the auto-regulatory effects of cell geometry alone. We argue that one such form of control could arise from the differential properties of newly created cell faces. These faces are developmentally distinct through their genesis, which could also influence their local biochemical state. Another possible mechanism could involve polarization effects induced by localization of Auxin transporters, such as the PIN family of protein^55,56^. In either case the local biochemical state could impact MT dynamics, such as to impact their stability. Here, we limit ourselves to local enhancement of MT stability in the basal cell face, achieved by 2-fold increase of the local catastrophe rate *r_c_* in the face other than the basal face. This parameterization increased the average innate MT length to 3.5 *μm* in that face, where the average MT length in the basal face was 4.5 *μm*. As panel C in Fig 16 shows, we again find the two major orientations found in the default, but now with slightly decreased propensities (cluster 1 ≈ 37% and cluster 2 ≈ 49%). However, a new orientation appears in ≈ 14% of realizations, in an unexpected tilted direction. This shows that face stability indeed provides an additional input for independent control, that is able to (at least partially) override the two geometrical mechanisms based on avoidance of self-intersections and catastrophe-inducing edges.

## Discussion

The modelling framework described in this article allows the simulation of cortical MT dynamics on triangulated approximations of essentially arbitrary three dimensional shapes, allowing for the first time to address the interplay among cell shape and MT array organisation on realistic cell shapes, which recent work has highlighted as being important^4^.

A crucial ingredient of the system is the ability to deal with localized variations in the MT dynamics. The well established propensity of MTs to undergo catastrophes when crossing high curvature regions between distinct cell faces is readily implemented, by identifying —for now by hand, but potentially automatically in the future — the regions involved. The second, is the, for now putative, yet biologically reasonable assumption of distinct levels of MT stability on different cell faces, due to the difference in their development, e.g. newly created faces due to cell division versus cell faces that expand through growth. The results, both on the simplified cuboidal geometry and the complex leaf pavement cells of *Nicotiana benthamiana* and *Hedera helix*, show that cell geometry alone already strongly restricts the possible array orientations. In the leaf pavement cells it can, moreover, predict the occurrence of non-trivial features such as band formation around neck-like protrusions. Both edge-catastrophe and face stability effects can then serve to fine-tune, or in some cases override, the spectrum of array orientations already limited by the cell geometry per se. Heuristically, the interplay between these different effects can be summarized as follows: (i) geometry favors MT trajectories that collectively form closed geodetic paths on the surface (ii) edge-catastrophes will select those closed paths that have fewest edge crossings, while (iii) enhanced face stability will select those paths that maximally intersect the selected face. It is precisely the balance between the latter two effects, which depending on the specific geometry can be either synergistic or antagonistic, that could provide a mechanism of cellular control over array orientation. As a rule, interphase array orientation parallels the orientation of the pre-prophase band, which in turn is a predictor for the location and orientation of the future division plane^51,52^. This suggests that insights in MT organisation from our simulations could be used to address long-standing issues in plant morphogenesis, which to a large degree is governed by the interplay between division plane orientation and cell growth. A first hint in that direction comes from our result on the reconstructing the possible division planes in merged daughter cells of *Hedera helix.*

Here we did not dwell on additional relevant factors such as the role of anisotropic MT-bound nucleation^57,58^ and MT severing by Katanin^59^. Both these effects, however, have already been implemented in the underlying MT interaction algorithms, and have been reported on elsewhere^60,61^.

Recent studies have provided evidence that MT dynamics is sensitive to the distribution of mechanical stress in the cell wall^62,63^. Our modelling framework has the ability to encode domain specific parametrization of MT dynamics. Therefore, once the precise influence of mechanical stress on the dynamics of MT is known, our modelling framework should allow the effects of a static stress distribution to be incorporated. In the current form, however, our modelling framework relies on the static template of a triangulated surface. It can therefore arguably only straightforwardly be applied to non- or slowly growing plant cells, for example those in early embryo development, where cell shape evolve slowly compared to e.g. division time.

In the context of growing cells, such as anisotropically growing root epidermal cells, the feedback mechanism between cortical MT dynamics, cell shape evolution and growth induced anisotropic stress needs to be considered. It will be a challenging, but potentially feasible task to extend our framework to simulate MT dynamics on growing cells as well, where cell growth e.g. is simulated by using the Finite Element Method^64–66^.

## Supporting Information

SI.1: Definition of edge-angles

SI.2: Implementation of edge-catastrophe

SI.3: Implemetation of MT stabilization

SI.4: Order parameter tensor

SI.5: Finite tubulin pool effect

SI.6: MT simulation on default cube surface

SI.7: Triangulation effect on MT-array orientation distribution

## Acknowledgements

This project was funded by an IPOP program grant of Wageningen University & Research. Computational work was carried out on the Dutch national e-infrastructure with support of the SURF Foundation.

## Author Contributions

BS and BM initiated the project. BC conceptualized the computational framework and developed the software. BM contributed technical improvements to the computational framework and the mathematical implementation of the order parameter. BC, BM and BS wrote the paper.

